## Supplementary material for "Using drivers and transmission pathways to identify SARS-like coronavirus spillover risk hotspots"

#### **Supplementary Materials**

**Muylaert el al.**

*Corresponding author email: r.delaramuylaert[at]massey.ac.nz

**This file includes:**

Figs. S1-S8

Table S1

Table S2

GIF 01

##### **Table S1. Average time to reach healthcare in areas where high emergent risk co-occurred with areas far from healthcare.**

| **Scenario** | **High-risk areas (N) far from healthcare** | **Average time to reach healthcare (hours)** | **min** | **max** | **SD** |
| --- | --- | --- | --- | --- | --- |
| Scenario 1 | 26 | 3.93 | 3.13 | 4.13 | 0.62 |
| Scenario 2 | 78 | 4.06 | 3.13 | 4.43 | 0.81 |
| Scenario 3 | 236 | 4.81 | 3.13 | 5.22 | 1.81 |
| Scenario 4 | 59 | 4.05 | 3.13 | 4.33 | 0.82 |

##### **Table S2. Hypothesized risk indicators informing the transmission scenarios, their rationale for inclusion, description and sources.** Original rasters were warped to 0.25 decimal degrees and World Geodetic System (WGS 84).

| **Higher-level indicator** | **Univariate spatial layers** | **Rationale for inclusion** | **References** | **Spatial layer details** | **Spatial layer source** |
| --- | --- | --- | --- | --- | --- |
| Landscape change (all scenarios) | Three layers were used, representing anthropogenic stressor intensities of: Built up area; Energy and mining; Agriculture and harvest. | Coronavirus shedding may be higher in human-dominated areas. Mining and agricultural areas are a signal of human activity even when population counts are low and can represent the margins where natural host habitat may be closer to human encounters. | (Anthony et al. 2017) | Summarizes land use intensity by human modification in 2017 (~1km). | (Theobald et al. 2020) |
| Landscape change (all scenarios) | Forest quality. | Emerging infectious disease risk is elevated in forested tropical regions experiencing land-use changes and where wildlife biodiversity (mammal species richness) is high. | (Allen et al. 2017) | Forest landscape integrity index, where highest values indicate highest quality (low=0, high=10) for 2019 (~1 km). It is based on inferred and observed human pressures (infrastructure, agriculture, tree cover loss) and loss of forest connectivity. | (Grantham et al. 2020) |
| Landscape change (all scenarios) | Risk of cover loss based on threats and dynamics. | Theory on land-use induced spillover; Agricultural land-uses exacerbate many infectious diseases in Southeast Asia (malaria, Schistosomiasis, Spotted fever, hookworms). | (Shah et al. 2019; Plowright et al. 2021; Rulli et al. 2021) | It informs the risk of a forest becoming removed in the future (transition potential, ~1 km), based on neural network models using historical data (2001-2014) from low (0) to high risk (1). Here we use continental model outcomes and not global, as the regional model estimates for Asia had better performance than the global model.. | (Hewson et al. 2019)  <https://futureclimates.conservation.org/riskstreecoverloss.html> |
| Potential secondary host (Scenario 2, Scenario 4) | Pigs. | Coronaviruses with origins tracing to bats causing disease in pigs. Sporadic infections cannot be excluded, but large-scale SARS-CoV-2 transmission among pigs is unlikely  (Sikkema et al. 2022). Respiratory illness symptoms have been associated with human contact with wildlife and livestock (Li et al. 2019). | (Zhou et al. 2018; Sikkema et al. 2022) | Areal-weighted GLW model (’Aw.tif’ files) from GLW3 Gilbert's livestock of the world estimates for 2010 (~10 km). This layer's original data spreads individuals of a census polygon evenly, so the density of animals in each pixel corresponds to the average number of animals/km2 of suitable land in the census unit. | (Gilbert et al. 2018) |
| Potential secondary host (Scenario 2, Scenario 4) | Cattle, bovid livestock. | Recent evidence from Germany. Concerns on anthropozoonotic infections of cattle reported as the presence of a preexisting coronavirus did not protect from infection with another betacoronavirus in a study. Also, multiple infections of individual animals might lead to recombination events between a SARS-like coronavirus and Bovine Coronavirus, a phenomenon already described for other pandemic coronaviruses. | (Wernike et al. 2022; Ulrich et al. 2020) | Areal-weighted GLW model (’Aw.tif’ files) from Gilbert's livestock of the world estimates for 2010 (~10 km).  This layer's original data spreads individuals of a census polygon evenly, so the density of animals in each pixel corresponds to the average number of animals/km2 of suitable land in the census unit. Results with all bovid livestock in the supplements (buffalo, cattle, goat, sheep). | (Gilbert et al. 2018) |
| Potential secondary host (Scenario 3, Scenario 4) | Wild mammals minus known bat hosts. | EID risk is elevated in forested tropical regions experiencing land-use changes and where wildlife biodiversity (mammal species richness) is high. SARS-Cov-2 has been detected in wildlife (spillback events). | (Nerpel et al. 2022; Allen et al. 2017) | IUCN data (~30 km), Search on 2022-04-04. Original Mollweide projection was warped to WGS84 in QGIS 3.24 after subtracting known bat host ranges. | <https://www.iucnredlist.org/resources/other-spatial-downloads#SR_2021_3> |
| Primary host (all scenarios) | Average estimated number of species of known bat hosts. | Peak of sarbecovirus hosts in Asia; Both the evolutionary and ecological aspects of emergence risk are higher in southeast Asia—a fact that will only become more relevant, as bats track shifting climates and exchange viruses with other species, creating a hotspot of elevated cross-species transmission unique to the region. Experimental evidence for bat (SARS-like) coronaviruses viruses infecting human cells, | (Muylaert et al. 2022; Sánchez et al. 2022; Forero et al. 2022; Temmam et al. 2022) | Average values used from the two sources. Sánchez et al. (2022) data (~1 km areas of habitat) was resampled to match Muylaert et al. (2022) resolution (0.25 dd). | (Muylaert et al. 2022; Sánchez et al. 2022) |
| Exposure (all scenarios) | Human population counts | Population size is a crucial transmissibility factor for SARS-like disease spread. | Sam John et al in prep  (Zhang et al. 2021) | Worlpop unconstrained global mosaics of population counts for 2020 (~1 km native resolution). | <https://hub.worldpop.org/geodata/listing?id=64> |
| Detection and spread in humans (not in the Scenarios) | Travel time to healthcare | City remoteness and hence access to healthcare are key to understanding zoonotic disease outbreaks. They can be used to understand early detection and connectivity. | (Winck et al. 2022) | Travel time to healthcare (motorized minutes, 1 km). This layer provides travel times to a nearest geolocated hospital or clinic. Hospital and clinic definition varies among countries but they assume they are: Fixed facilities providing urgent or emergency medical care with an entry subtype indicating they were a hospital or clinic, that were open in August 2019. Mobile or temporary clinics for providing healthcare in remote areas are not taken into account. Data completeness for healthcare varies by country: China (53451) has the largest number of detected facilities in the world, followed by other Asian countries considered in our analysis, such as India, Indonesia, Thailand, Malaysia, Sri Lanka, etc.. So we assume that there is good coverage in Asia, and according to the authors Google had the best data sources for Asian countries. The Philippines had 2358 facilities detected (1 km pixel resolution). | (Weiss et al. 2020) |

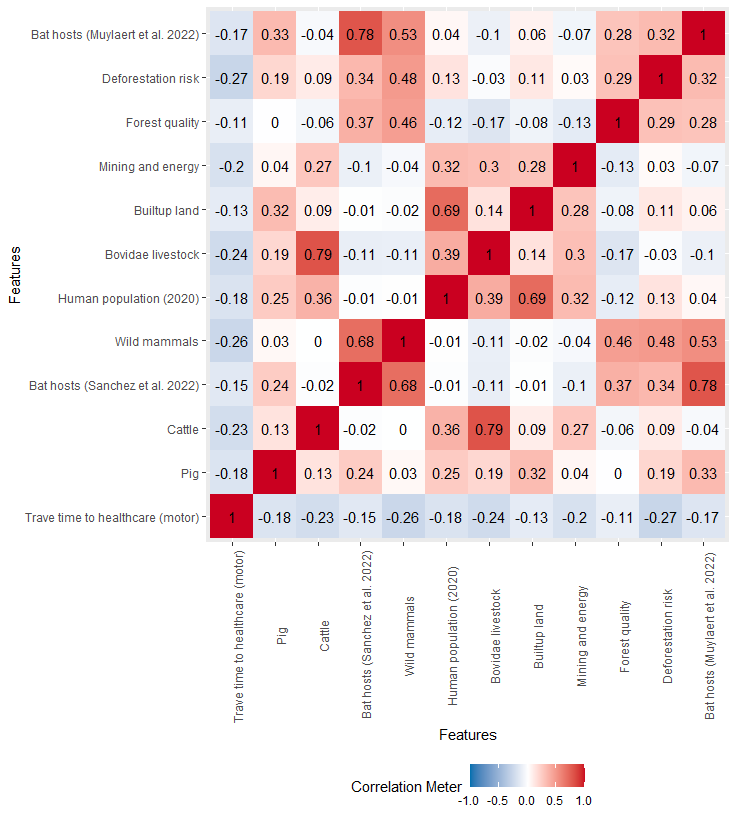

##### **Figure S1. Product-moment correlation values (*r*) of selected variables.** Known bat hosts were combined in a single layer after averaging their values. Results with the cattle-only version are displayed in the main text, and Bovidae livestock in the supplements.

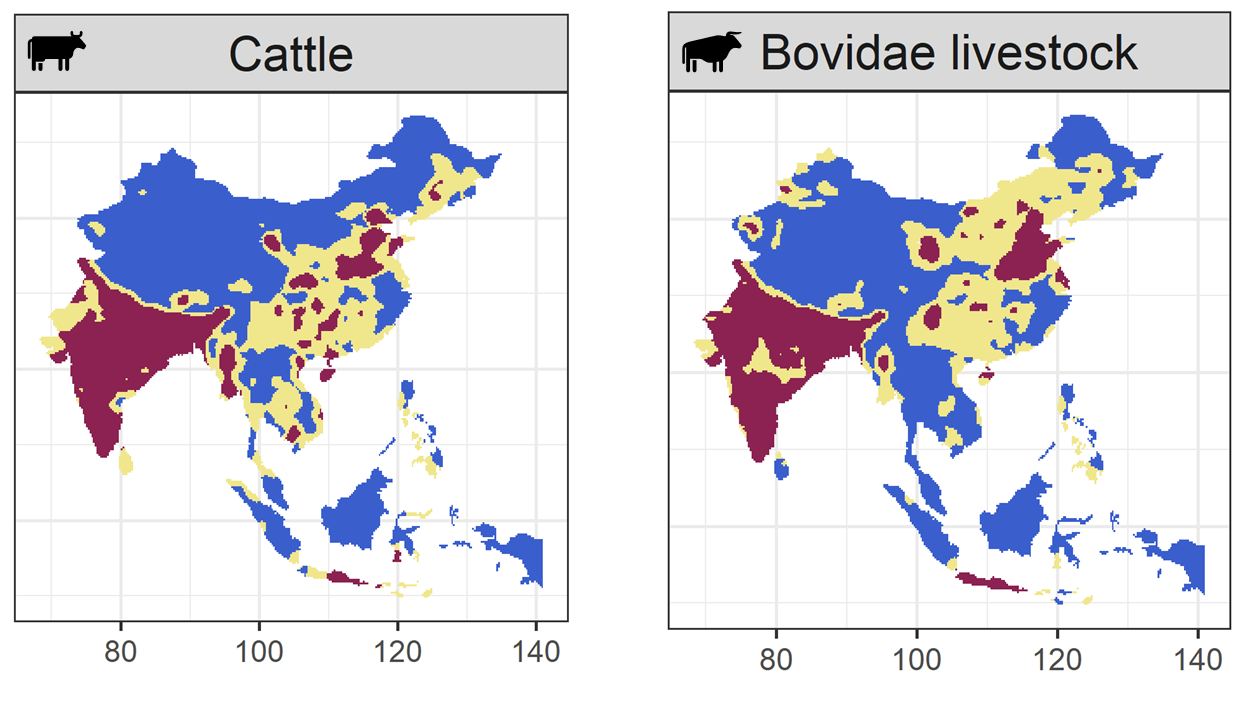

##### **Figure S2. Hotspot values for cattle and all Bovidae livestock.** Hotspots in dark red, intermediate zones in yellow, coldspots in blue.

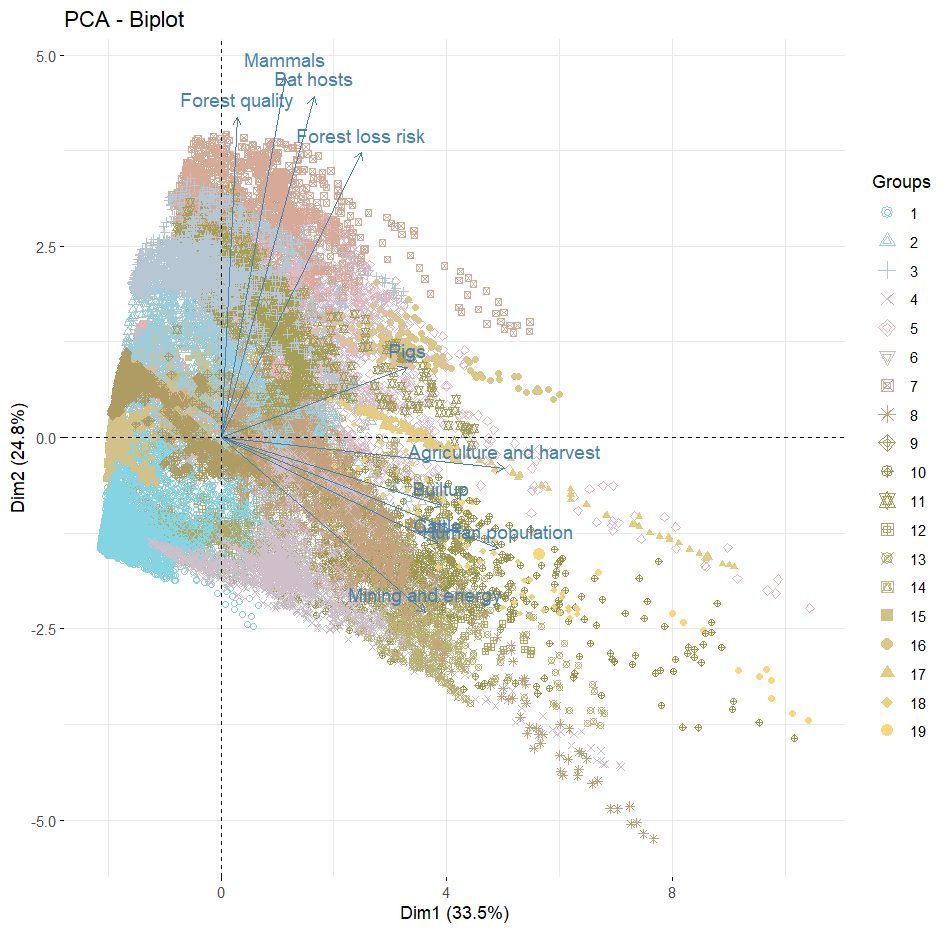

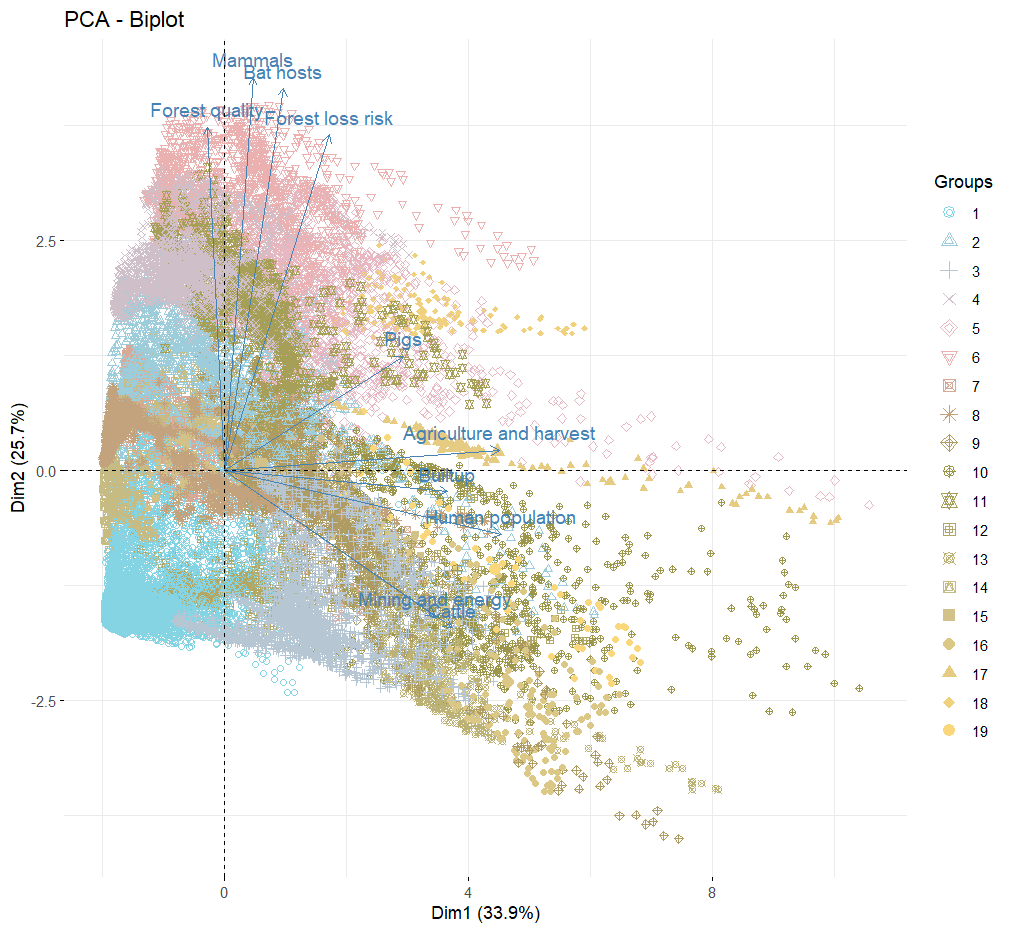

**Figure S3. Principal component analysis (PCA) biplot indicates variation between 19 clusters defined by multivariate spatial cluster analyses considering all variables (scenario 4).** Upper panel: cattle-only version. Bottom panel: Bovidae livestock version.

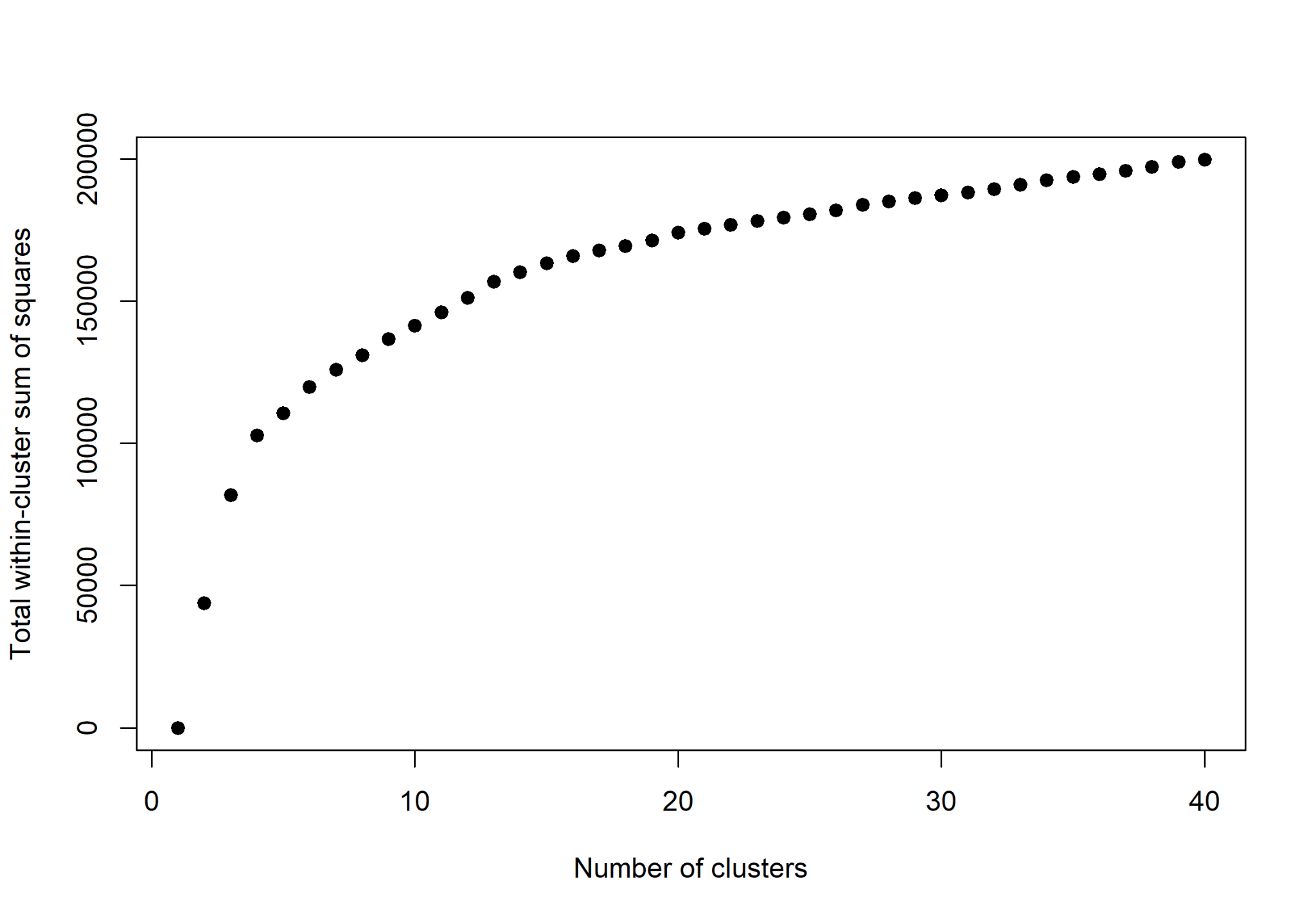

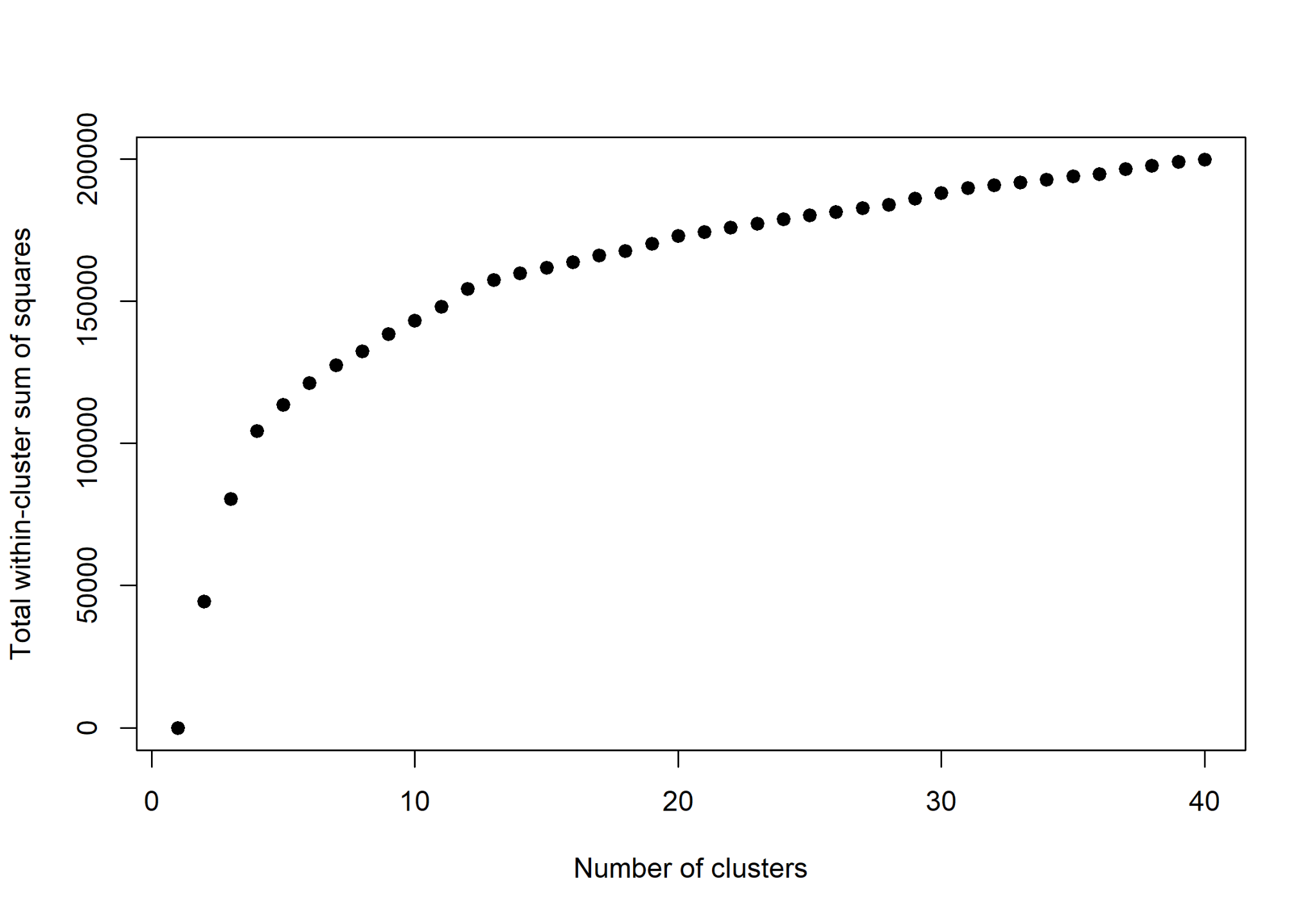

##### **Figure S4.** Skater within-cluster sum of squares variation from 1 to 40 clusters for all selected variables (scenario 4). The optimal number of clusters informed by the max-p algorithm was 9 and 19 (respectively, for 10% and 5% human population used as minimum bound variables). Upper panel: Cattle-only version. Bottom panel: Bovidae livestock version.

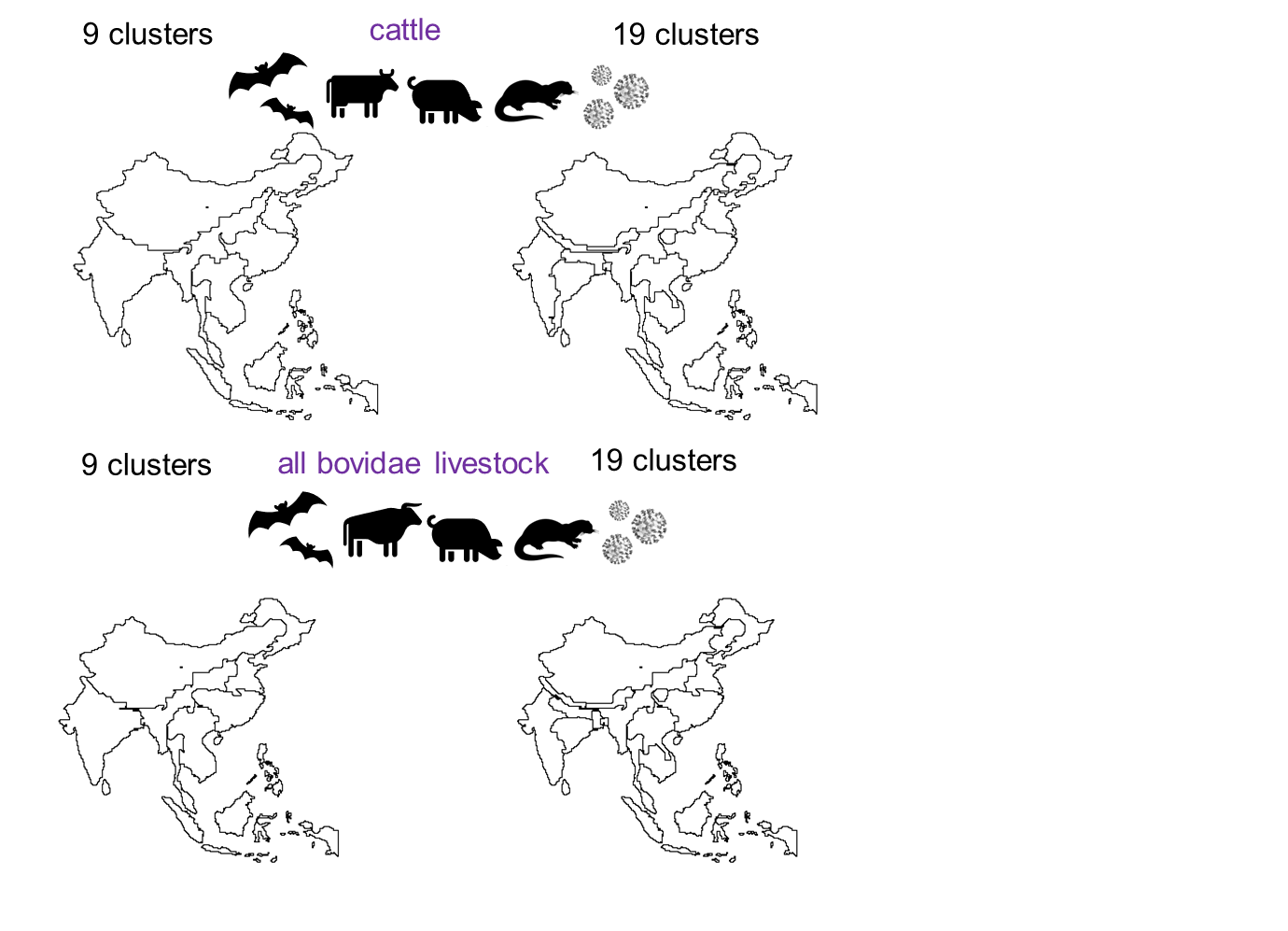

##### **Figure S5. Hierarchical nature of the spatial clusters with 9 and 19 optimal number of clusters considering the global scenario (scenario 4).** Results presented with 19 clusters are in the main text. Upper panel: Cattle-Only. Bottom: Bovidae livestock.

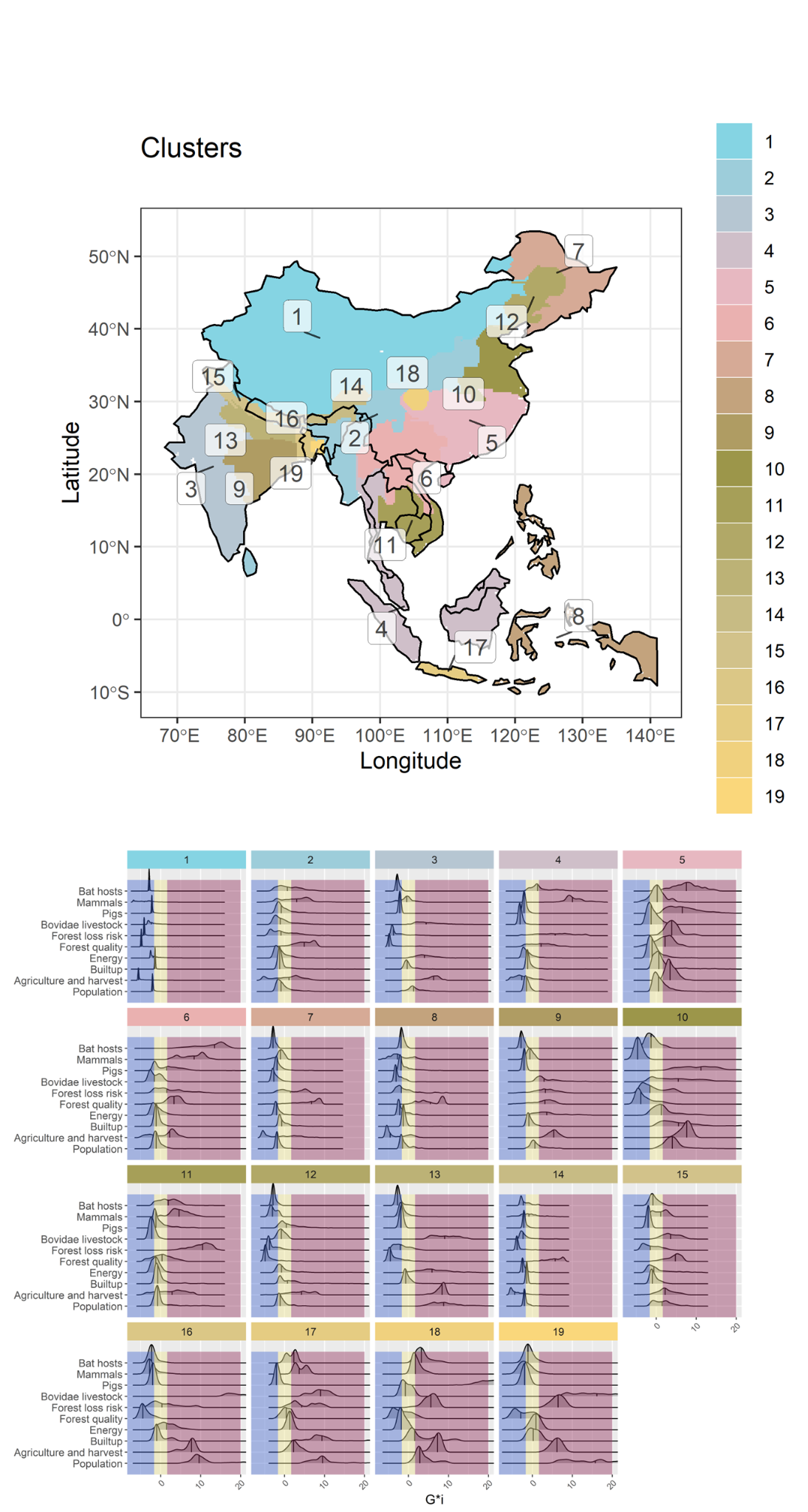

##### **Figure S6. Optimal number of multivariate clusters of all selected components associated with potentially new emerging SARS-like coronavirus (scenario 4).** This version uses Bovidae livestock instead of cattle-only.

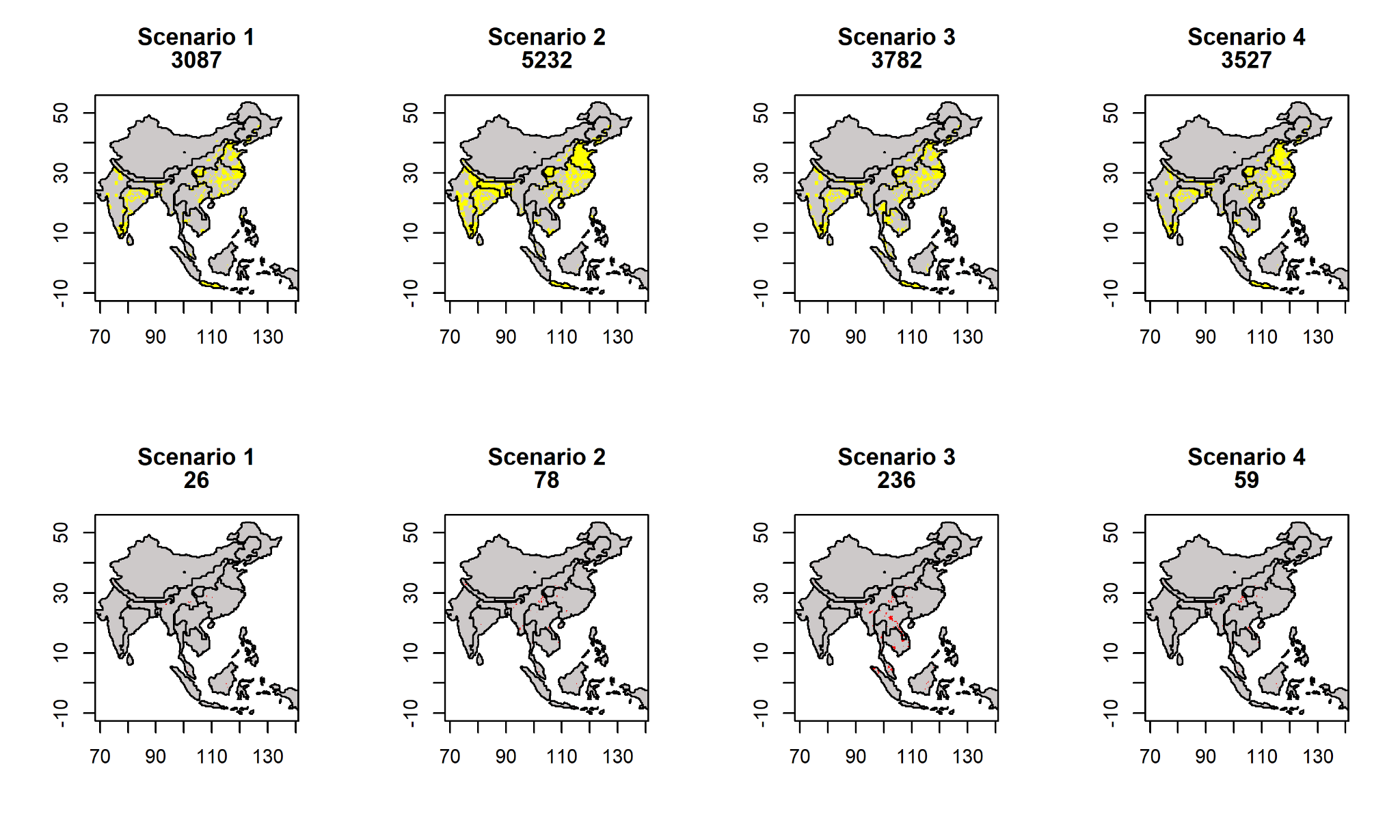

**Figure S7. Risk associated with transmission scenarios according to time to reach healthcare (lower and higher quantiles for healthcare access).** Boundaries in black represent the 19 clusters. Upper panel shows areas that are close from healthcare, with high hotspot overlap, in yellow. Bottom panel shows areas that are far from healthcare, with high hotspot overlap, in red. The number below every title corresponds to the grid count for the colour value. Landscape, human population and known bat hosts are included in all models, and are the sole indicators in Scenario 1, representing direct transmission. To incorporate indirect transmission through secondary hosts, mammalian livestock are included in Scenario 2, wild mammals in Scenario 3, and both mammalian livestock and wild mammals in Scenario 4.

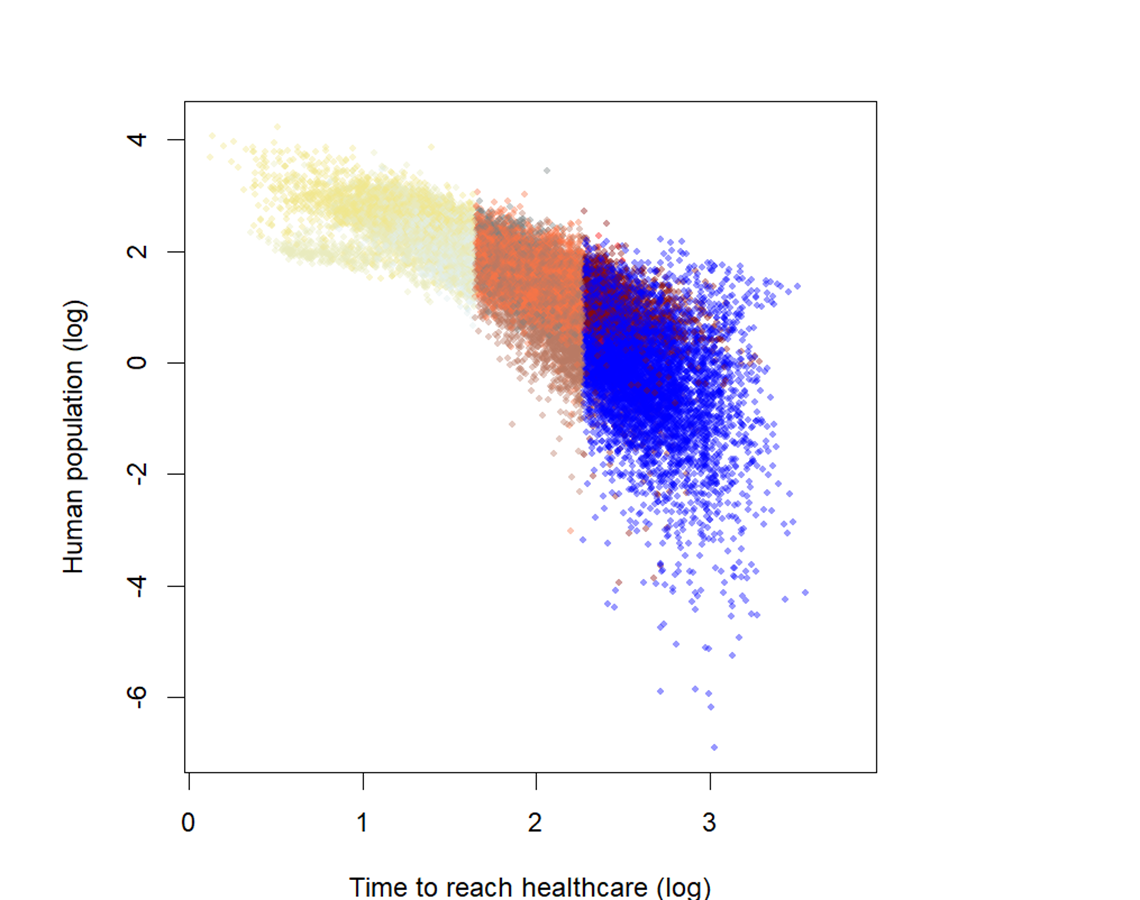

##### **Figure S8.** Human population variation according to motorized travel time and coloured by quantiles from the bivariate map of inferred risk from Scenario 4 as a function of time to reach healthcare.

**GIF 01. Comparison among different scenarios of risk and access to health care.** The two data layers were divided in three quantiles and 9 unique legend color codes.

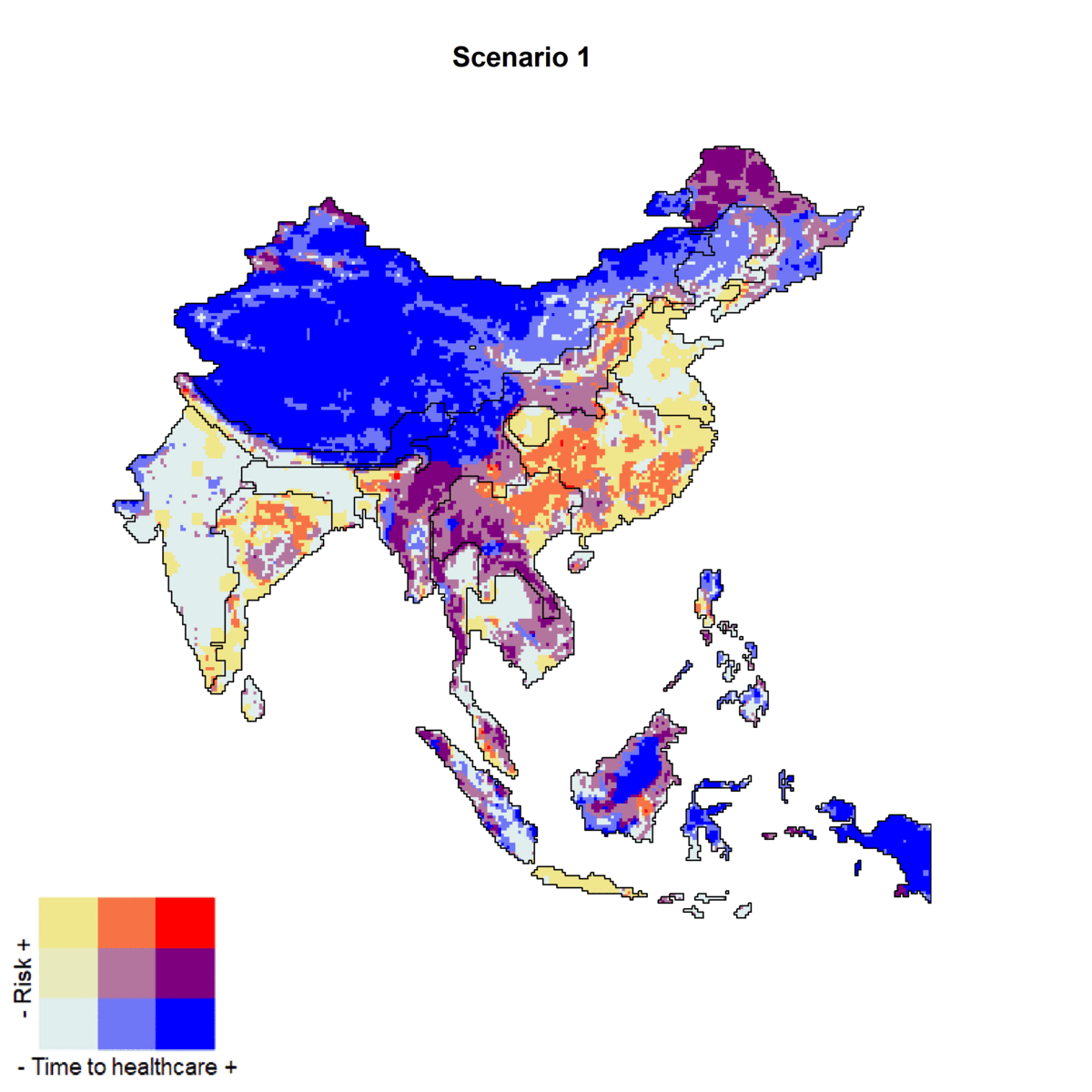
